# SARS-CoV-2 entry and fusion are independent of ACE2 localization to lipid rafts

**DOI:** 10.1101/2024.07.13.603361

**Authors:** William Bolland, Inès Marechal, Chloé Petiot, Françoise Porrot, Florence Guivel-Benhassine, Nicoletta Casartelli, Olivier Schwartz, Julian Buchrieser

## Abstract

Membrane fusion occurs at the early stages of SARS-CoV-2 replication, during entry of the virus, and later during the formation of multinucleated cells called syncytia. Fusion is mediated by the binding of the viral Spike protein to its receptor ACE2. Lipid rafts are dynamic nanodomains enriched in cholesterol and sphingolipids. Rafts can act as platforms for entry of dìerent viruses by localizing virus receptors, and attachment factors to the same membrane microdomains. Here, we first demonstrate that cholesterol depletion by methyl-beta-cyclodextrin inhibits Spike mediated fusion and entry. To further study the role of ACE2 lipid raft localization in SARS-CoV-2 fusion and entry, we design a GPI-anchored ACE2 construct. Both ACE2 and ACE2-GPI proteins are similarly expressed at the plasma membrane. Through membrane flotation assays, we show that in dìerent cell lines, ACE2-GPI localises predominantly to raft domains of the plasma membrane while ACE2 is non-raft associated. We then compare the ability of ACE2 and ACE2-GPI to permit SARS-CoV-2 pseudovirus entry and syncytia formation and replication of dìerent viral variants. We find little dìerence in the two proteins. Our results demonstrate that SARS-CoV-2 entry and fusion are a cholesterol dependent and raft-independent process.

**IMPORTANCE:** Rafts are often exploited by viruses and used as platforms to enhance their entry into the cell or spread from cell-to-cell. The membrane localization of ACE2 and the role of lipid rafts in SARS-CoV-2 entry and cell-to-cell spread is poorly understood. The function of lipid rafts in viral fusion is often studied through their disruption by cholesterol-depleting agents. However, this process may have ò-target impacts on viral fusion independently of lipid-raft disruption. Therefore, we created an ACE2 construct that localizes to lipid rafts using a GPI anchor. Conversely, wild-type ACE2 was non-raft associated. We find that the localization of ACE2 to lipid rafts does not modify the fusion dynamics of SARS-CoV-2.

## INTRODUCTION

Enveloped viruses, such as coronaviruses, use fusogens to facilitate entry and infection at the cell surface or within the endocytic pathway (Dimitrov, 2004; Jackson et al., 2022). SARS-CoV-2 Spike protein binds to the angiotensin converting enzyme 2 (ACE2) receptor, leading to conformational changes in Spike to allow release of the fusion peptide through cleavage of the S2’ site by cell surface proteases, such as TMPRSS2, or endosomal proteases such as cathepsins (Bestle et al., 2020; Jaimes et al., 2020). During the late stages of viral replication, Spike leaks from the viral packing site in the endoplasmic reticulum and Golgi network to the cell surface due to a suboptimal COPI binding motif (Cattin-Ortolá et al., 2021). There, Spike interacts with neighbouring ACE2-expressing cells, leading to cell-to-cell fusion and the formation of multinucleated cells, known as syncytia. Syncytia formation by SARS-CoV-2 has been well documented in *in vitro* studies and from analyses of lung samples of COVID-19 patients (Braga et al., 2021; Buchrieser et al., 2020; Bussani et al., 2020, 2023). We and others have shown that the SARS-CoV-2 variants possess dìerent syncytia-forming capacities due to mutations in the Spike protein (Bolland et al., 2024; Meng et al., 2022; Planas et al., 2024; Rajah et al., 2021; Saito et al., 2022) and that high fusogenic activity is linked to increased cytopathy (Bolland et al., 2024). The role of syncytia during SARS-CoV-2 infection includes dissemination of the virus between cells while escaping the humoral immunity (Rajah et al., 2022; Zeng et al., 2022). However, questions remain surrounding the role of syncytia in pathogenesis and the mechanisms regulating syncytia formation.

Lipid rafts are highly dynamic nanodomains (< 200 nm) present within the plasma membranes of all eukaryotic cells, that are enriched in cholesterol and sphingolipids. Post-translational modifications of proteins, such as the addition of a glycosylphosphatidylinositol (GPI) anchor, localize the proteins to lipid rafts (Parkin et al., 2003). Rafts perform cellular functions such as signal transduction by concentrating and crosslinking signalling receptors, and exocytosis by organizing SNAP REceptor (SNARE) proteins at the plasma membrane (Chamberlain et al., 2001; Simons & Toomre, 2000).

Lipid rafts have been proposed to act as platforms for the entry of diverse enveloped and nonenveloped viruses through localization of the virus receptor, co-receptors, and attachment factors (Ripa et al., 2021). For example, human herpesvirus-6 recruits its receptor CD46 to rafts to permit infection (Tang et al., 2008), likewise with the attachment factors of hepatitis C virus (Kapadia et al., 2007), and the CCR5 and CXCR4 co-receptors of human immunodeficiency virus-1 (Cardaba et al., 2008; Kamiyama et al., 2009). Both SARS-CoV and SARS-CoV-2 have been proposed to employ ACE2 localization into rafts to facilitate host cell entry through the caveolin-dependent pathway (Palacios-Rápalo et al., 2021). However, the exact localization of ACE2 in the plasma membrane is disputed with reports suggesting it is a raft-associated protein and others suggesting the contrary (El Khoury & Naim, 2023; G. M. Li et al., 2007; X. Li et al., 2021; Warner et al., 2005). Therefore, clarification of the membrane localization of ACE2 and the importance of its localization in SARS-CoV-2 entry is needed.

The classical approach in studying the role of lipid rafts during viral replication is by their disruption by cholesterol-depleting agents such as cyclodextrins and statins (Sviridov et al., 2020; Zidovetzki & Levitan, 2007). Disruption of rafts in ACE2-expressing cells reduces entry of SARS-CoV and SARS-CoV-2, indicating that cholesterol plays an important role and suggested that lipid rafts may be involved in this process (El Khoury & Naim, 2023; X. Li et al., 2021; Y. Lu et al., 2008). Nevertheless, cholesterol depletion is not specific for rafts (Mahammad & Parmryd, 2008) and may have ò-target impacts on cell homeostasis, such as glucose transport, and membrane elasticity (Caliceti et al., 2012; Yang et al., 2016). Investigating the role of lipid rafts in SARS-CoV-2 fusion and entry through ACE2 modifications has not been explored.

Here, we confirm the importance of cholesterol in the entry and fusion of SARS-CoV-2 by depletion using Methyl β cyclodextrin (MβCD). We then show that wild-type ACE2 localizes to non-raft regions of the membrane and subsequently design a GPI-anchored ACE2 construct that localizes to lipid rafts. Through reporter assays for syncytia formation and pseudovirus entry, we find that ACE2 localization to lipid rafts has no significant impact on SARS-CoV-2 entry and fusion, except in HEK 293T cells where raft localization slightly increases syncytia formation and pseudovirus entry. We then corroborate these findings using SARS-CoV-2 virus by measuring viral replication and syncytia formation.

## METHODS

### Cell lines

HEK 293T, MRC5, U2OS, and HeLa cells and their derivatives were purchased from the ATCC or generously provided by fellow stà at Institut Pasteur. All cell lines were cultured at 37°C, 5% CO_2_ in DMEM media supplemented with 10% fetal bovine serum (FBS) and 1% penicillin/streptomycin antibiotics. For fusion assays, GFP-split stable cell lines were generated by lentivector transduction containing pQCXIP-derived plasmids for GFP1-10 or GFP11 subunits. GFP1-10 or GFP11 expressing cells were selected through puromycin resistance and cultured by supplementation of the media with puromycin (HEK 293T, U2OS, HeLa – 1 μg/mL; MRC5 – 10 μg/mL). HEK 293T and MRC5 ACE2 and ACE2-GPI stable cell lines were generated by lentivector transduction containing pLV-derived plasmids for ACE2 or ACE2-GPI. ACE2 and ACE2-GPI expressing cells were selected through hygromycin B resistance and cultured by supplementation of the media with 100μg/mL (HEK 293T) or 200 μg/mL (MRC5) hygromycin B. All cells used in this study were routinely checked and found negative for mycoplasma.

### SARS-CoV-2 isolates

The D614G and XBB.1.5 isolates were previously described (Planas et al., 2024). Briefly, the D614G and XBB.1.5 strains were isolated and cultured on VeroE6 or IGROV-1 cell lines respectively. The D614G strain (hCoV-19/France/GE1973/2020) was obtained from the National Reference Centre for Respiratory Viruses at Institut Pasteur. The XBB.1.5 strain was isolated from a nasopharyngeal swab of an anonymous individual at the Hopital Europeen Georges Pompidou (HEGP; Assistance Publique, Hôpitaux de Paris). Virus stock titers were calculated by TCID_50_ measurements as previously described. All work using SARS-CoV-2 virus was carried out in a BSL-3 laboratory, under the guidelines of the risk prevention service at Institut Pasteur.

### GFP-split fusion assay

SARS-CoV-2 Spike induced cell-to-cell fusion was measured using the GFP-split system. All transfections were carried out using lipofectamine 2000 reagent (ThermoFisher) and incubated on a shaking incubator at 1000 rpm 37°C for 2 hours. For HEK 293T, U2OS, and HeLa GFP-split expressing cell lines, the respective GFP1-10 cell lines were transfected with plasmids encoding ACE2 (pLV) or ACE2-GPI (pLV) plasmids in the presence or absence of TMPRSS2 (pcDEST). The GFP11 cell lines were transfected with plasmids encoding D614G Spik or XBB.1.5 Spike (pVAX1, Invitrogen). Control conditions were performed using a pQCXIP-Empty plasmid. Following transfection, cells were centrifuged at 500 g to pellet cells and the transfection mix was discarded. Cells were resuspended and GFP1-10 and GFP11 cells were co-cultured in µClear black 96-well plates (Greiner Bio-One) for 18 hours. Cells were cultured at the following densities (50:50 ratio of GFP1-10 and GFP11 cells): HEK 293T – 6.0 x 10^4^ cells per well; U2OS – 3.0 x10^4^ cells per well; HeLa – 5.0 x10^4^ cells per well. For MRC5 fusion assays, 2.5 x10^4^ MRC5 GFP1-10 cells stably expressing ACE2 or ACE2-GPI were co-cultured with 3.0 x10^4^ HEK 293T GFP11 cells transfected for D614G or XBB.1.5 Spike or a control plasmid. 18 hours post-transfection, fluorescence microscopy images of the cells were acquired using the Opera Phenix High-Content Screening System (PerkinElmer). Hoechst 33342 was added to the culture media at 1:10000 dilution to perform nuclei counting. GFP area and nuclei number were quantified using the Harmony High-Content Imaging and Analysis Software (PerkinElmer, HH17000012, v.5.0). Surface expression of ACE2, Spike, and TMPRSS2 was analysed as described in the “Flow cytometry” section.

For SARS-CoV-2 virus-induced cell-to-cell fusion assays, HEK 293T GFP1-10 and GFP11 cells were pooled and transfected with ACE2 or ACE2-GPI or a control plasmid for 24 hours. 6.0 x10^4^ HEK 293T cells were seeded per well in µClear black 96-well plates. MRC5 GFP1-10 and GFP11 cells stably expressing ACE2, or ACE2-GPI or control wild-type cells were seeded at a density of 5.0 x10^4^ cells per well. Cells were then infected with D614G or XBB.1.5 virus at MOI 0.1 for 48 hours. 48 hours post-infection, cell supernatant was collected for qRT-PCR analysis and cells were fixed with 4% paraformaldehyde (PFA), washed, and stained with Hoechst 33342. Intracellular Spike staining was performed with antibodies diluted in PBS + 1%-BSA, 0.1%-sodium azide, 0.05%-saponin (ThermoFisher) using mAb10 anti-S2 antibody (Planchais et al., 2022) as the primary and Alexa fluor 647-conjugated goat anti-human IgG as the secondary. GFP area and nuclei number were quantified on the Opera Phenix High-Content Screen System as described in the previous paragraph.

### SARS-CoV-2 pseudovirus entry

Pseudoviruses were produced by co-transfection of HEK 293T cells with plasmids encoding SARS-CoV-2 Spike (D614G or XBB.1.5), lentiviral proteins, and a luciferase reporter plasmid. Pseudoviruses were harvest 48 hours post-transfection and titres were quantified by measuring infectivity. The assessment of pseudovirus entry was carried out in HEK 293T cells transfected for 24 hours with ACE2 or ACE2-GPI or control plasmid and in MRC5 cells stably expressing ACE2 or ACE2-GPI or wild-type control cells. 4 x10^4^ HEK 293T cells or 3 x10^4^ MRC5 cells were seeded per well in white Cellstar 96 well cell culture plates (Greiner Bio-One). Pseudoviruses were added to the cell media at a 1:50 and 1:250 dilution. 48 hours after pseudovirus infection, 50 µL Bright-Glo substrate (Promega) was added to each well and incubated in the dark for 5 minutes prior to plate reading. Luciferase activity was measured using an EnSpire plate reader (PerkinElmer).

### MβCD drug treatment assays

The impact of MβCD on cell-to-cell fusion and SARS-CoV-2 pseudovirus entry was carried out in HEK 293T GFP-split cells and wild-type cells respectively. For cell-to-cell fusion assays, HEK 293T GFP1-10 cells were transfected with ACE2 or control plasmid, and GFP11 cells were transfected with D614G Spike or control plasmid for 24 hours. Cells were then treated with 2 mM MβCD or the equivalent volume of DMSO for 3 hours on a shaking incubator at 1000 rpm 37°C. Cells were then pelleted, washed, and seeded at 6.0 x10^4^ cells per well (50:50 ratio of both GFP1-10 and GFP11 cells). GFP quantification was carried out 3 hours after seeding on the Opera Phenix High-Content Screen System. For pseudovirus entry, HEK 293T cells were transfected with ACE2 or control plasmid for 24 hours. Cells and D614G Spike pseudovirus were treated with 2 mM MβCD or the equivalent volume of DMSO for 3 hours, cells on a shaking incubator at 1000 rpm 37°C. Cells were pelleted, washed, and seeded. Cells were infected with 1:50 pseudovirus and luciferase activity was measured as described in the earlier “SARS-CoV-2 pseudovirus entry” section.

### Membrane flotation assay and western blotting

For flotation assays, 1.5 x 10^7^ cells were cultured and lysed per condition. Cells were lifted and resuspended in 300 μL NTE bùer [50 mM Tris-HCL (pH 7.4), 50 mM NaCl, 5 mM EDTA] to wash. Cells were then centrifuged at 500 g and resuspended in NTE bùer + 1% Triton X-100 + Roche complete protease inhibitor on ice for 1 hour to lyse cells. Cell lysates were then transferred to a Dounce homogenizer and passed 10 times with the pestle avoiding bubble formation. Lysates were transferred to tubes and centrifuged at 2500 g, 4°C for 10 minutes to remove cell debris and nuclei. Lysates were gently mixed with 2 mL of 60% OptiPrep iodixanol density gradient medium (Sigma-Aldrich) within 15 mL ultracentrifuge tubes. A density gradient was then created by slowly adding 6 mL of 30% OptiPrep and then 2.5 mL 5% OptiPrep diluted in NTE bùer. Samples were loaded into an ultracentrifuge and spun at 100,000 g for 18 hours. Samples were then aliquoted gently into fractions and stored at -20°C prior to western blot analysis.

For western blot analysis, 10 μL of fraction was loaded per well. Samples were reduced using Laemmli sample bùer 4x (Bio-Rad) and NuPAGE Sample reducing agent 10x (ThermoFisher) for 5 minutes at 95°C. NuPAGE 4-12% Bis-Tris Gels (Invitrogen) were used to run samples. PageRuler prestained protein ladder (ThermoFisher) was used as a kDa reference. Following the transfer, membranes were blocked in PBS + 5%-BSA overnight at 4°C. Antibodies were incubated for 1 hour at room temperature and diluted in PBS + 1%-BSA, 0.05%-Tween, 0.1%-sodium azide. Primary antibodies used were for ACE2 - goat anti-hACE2 (R&D systems, 1 μg/mL) and for flotillin-1 - rabbit anti-flotallin-1 (ThermoFisher, 1 μg/mL). Membranes were washed three times in PBS + 0.05%-Tween between each antibody incubation. Secondary antibodies used were anti-goat IgG DyLight 680 (Life technologies, 1:5000) and anti-rabbit IgG DyLight 800 (Invitrogen, 1:5000). Membranes were revealed using the Licor imager and analyzed using Image Studio Lite v5.2.5 software.

### Flow cytometry

Surface staining was performed on live cells and all primary and secondary antibodies were diluted in MACS bùer [PBS + 0.5%-BSA, EDTA 2 mM] and incubated at 4°C for 30 min. Primary antibodies used were: for ACE2 - 1 μg/mL anti-ACE2 VHH-B07-Fc (Bretol el al., manuscript in preparation), for Spike - 1 μg/mL LY-CoV404 (Bruel et al., 2022), for TMPRSS2 – anti-TMPRSS2 VHH-A01-Fc (Saunders et al., 2023). Cells were washed in 100 μL PBS between primary and secondary antibody incubations. Alexa fluor 647-conjugated goat anti-human IgG was used as the secondary antibody (Invitrogen, A-21445, 1:500). For phospholipase C (PLC) treatment, cells were incubated with 100 or 200 U/μL PLC for 2 hours at 37°C on a shaking incubator at 1000rpm. Cells were washed and resuspended before analysis of ACE2 surface levels as described above. For soluble SARS-CoV-2 spike RBD binding analysis, cells were stained with 4.5 μg/mL soluble Wuhan RBD-biotinylated (Planchais et al., 2022) at 4°C for 30 min. Cells were washed in 100 μL PBS and then Alexa fluor 488-conjugated streptavidin (ThermoFisher, S11223, 1:500) was incubated at indicated timepoints at 4°C for 30 min. All cells were fixed in 4% PFA and resuspended in 200 μL PBS before acquisition on the Attune NtX flow cytometer (ThermoFisher). Data were analyzed using FlowJo software (BDBioSciences).

### qRT-PCR

For quantification of viral RNA release, cell supernatants were collected and inactivated by diluting 1:4 in H_2_O and incubating at 80°C for 20 min. For amplification SARS-CoV-2 N-gene forward (5’-TTACAAACATTGGCCGCAAA-3’) and reverse (5’-GCGCGACATTCCGAAGAA -3’) primers were used at 10 μM (X. Lu et al., 2020). Luna One-step qRT-PCR kit (New England Biolabs) was used with 1 μL supernatant. A standard curve was generated by 1:10 serial dilution of EURM-019 ssRNA SARS-CoV-2 fragments (European Commission). Analysis was performed with the QuantStudio 6 Flex Real-Time PCR machine (ThermoFisher).

### Statistical analysis

Calculations were performed using Excel 365 (Microsoft). Figures and statistical analyses were conducted using GraphPad Prism 9. Statistical significance between dìerent groups was calculated using the tests indicated in each figure legend. No statistical methods were used to predetermine sample size.

## RESULTS

### Cholesterol depletion reduces SARS-CoV-2 pseudovirus entry and Spike induced cell-to-cell fusion

We first investigated if cholesterol was required in the membrane of the virus or in the membrane of the cell expressing ACE2. To investigate this in SARS-CoV-2 entry, we incubated ACE2 expressing HEK 293T cells and SARS-CoV-2 pseudovirus with 2 mM MýCD prior to infection. Following cholesterol depletion in the ACE2 expressing target cells, the mean RLU decreased significantly by 50% while treatment of the pseudovirus had no significant àect on entry (Fig. 1A). Next, we investigated the role of cholesterol in Spike-ACE2 mediated cell-cell fusion, we used the GFP-split system (Buchrieser et al., 2019, 2020; Planas et al., 2021) as a reporter of fusion. We treated GFP11 cells expressing Spike and GFP1-10 cells expressing ACE2 with 2 mM MβCD prior to co-culturing of the cells. GFP quantification showed a significant 45% and 47% reduction in GFP area when either Spike or ACE2 expressing cells are depleted for cholesterol, respectively, and a cumulative 76% decrease when both cells are treated (Fig. 1B). Taken together, cholesterol within the cell plasma membrane is required to allow entry of pseudovirus as well as fusion of the cells leading to syncytia formation.

**Figure 1.**
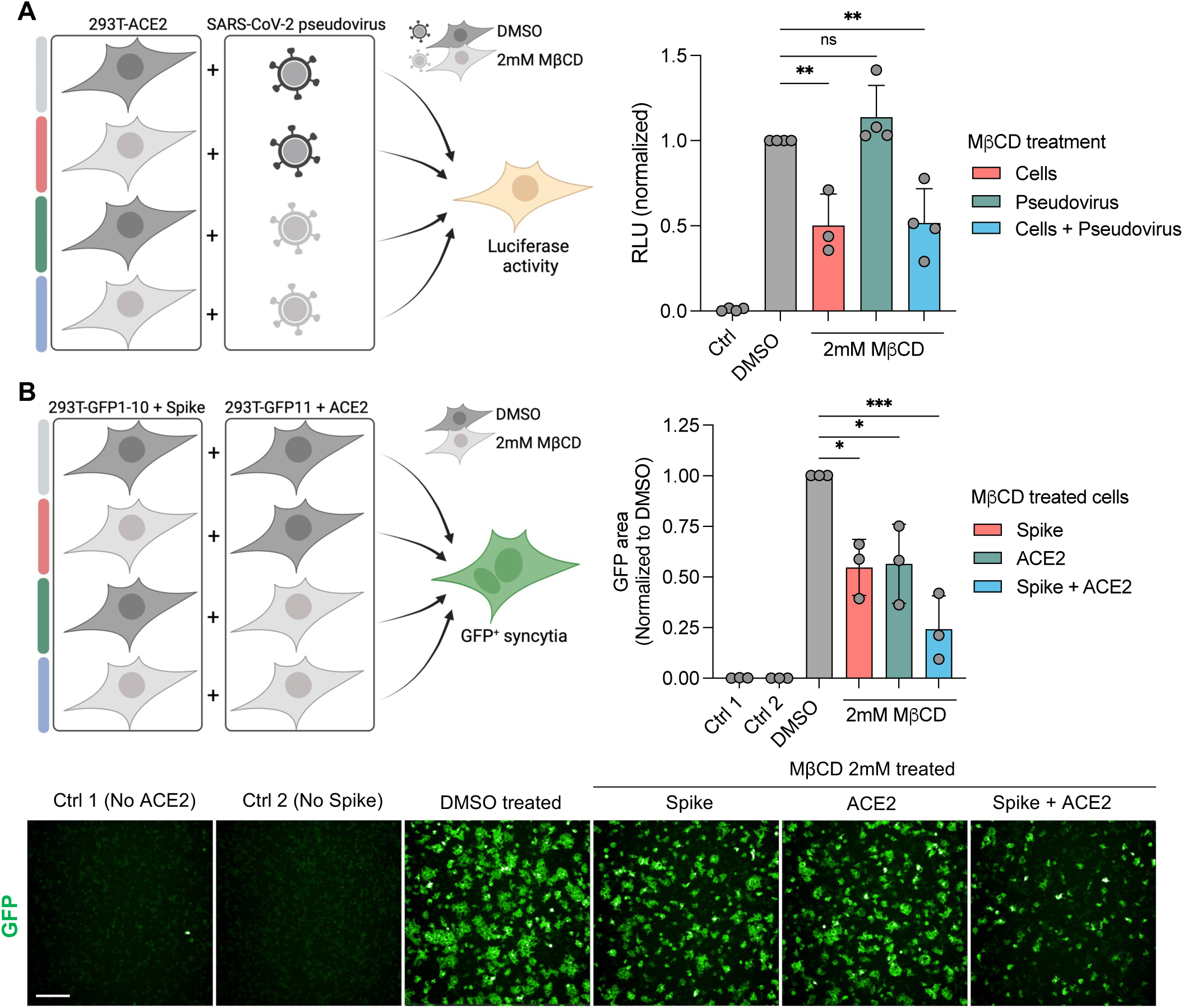
Plasma membrane cholesterol depletion reduces SARS-CoV-2 fusion and entry. **(A)** 293T GFP-split cells were transfected with SARS-CoV-2 Spike (GFP 1-10 expressing cells) or ACE2 (GFP 11 expressing cells) 24h before MβCD or DMSO treatment. Treatment of cells with 2 mM MβCD or DMSO control was carried out prior to co-culturing of cells as shown (Left). GFP area was quantified 3h after co-culture for a readout of cell fusion. Data were normalized to the Spike+ACE2 DMSO condition. Data are means ±s.d. of three independent experiments. **(B)** 293T cells were transfected with ACE2 24h before pseudovirus infection. Treatment of pseudovirus with 2 mM MβCD or DMSO control was carried out for 3h prior to addition of the pseudovirus. Cells were infected for 48h prior to addition of luciferase substrate and luminescence quantification. Ctrl 1 = No ACE2; Ctrl 2 = No Spike. Data were normalized to the ACE2 DMSO condition. Data are means ±s.d. of three independent experiments. Below: Immunofluorescence images of 293T GFP-split cells taken from one of three independent experiments. Scale bar: 200 μm. For A, and B, Ordinary One-Way ANOVA tests were performed with Tukey’s multiple comparison test to compare to DMSO treatment, \**P* < 0.05, \*\**P* < 0.001, \*\*\**P* < 0.0001, ns = not significant.

### GPI-anchored ACE2 localizes to lipid rafts while wild-type ACE2 is non-raft associated

Cholesterol depleting agents impact a variety of cellular processes and do not specifically act on lipid rafts (Caliceti et al., 2012; Mahammad & Parmryd, 2008). Therefore, to investigate the role of lipid rafts during SARS-CoV-2 entry and fusion without cholesterol depletion, we designed an ACE2 construct in which the transmembrane domain (TD) and CT were exchanged for a Thy-1 GPI-anchored protein (Herein referred to as ACE2-GPI; Fig. 2A). We asked whether the construct was similarly expressed on the cell surface compared to wild-type ACE2., HEK 293T cells were transfected with a serial dilution of either ACE2 or ACE2-GPI plasmid and expression was examined by surface staining with an anti-ACE2 antibody. Analysis by flow cytometry revealed that both the percentage of ACE2-positive cells and the mean fluorescence intensity (MFI) of the cells showed no significant dìerences between ACE2 and ACE2-GPI surface levels (Fig. 2B). We next investigated the rate of endocytosis of ACE2-GPI compared to ACE2. HEK 293T cells, expressing either protein, were incubated with SARS-CoV-2 biotinylated-RBD, washed, and incubated with streptavidin-AF-488 at the indicated timepoints. Over the course of 24h both ACE2 and ACE2-GPI surface level decreased at the same rate, indicating both constructs display similar surface turnover (Fig. 2C). To further validate the ACE2-GPI construct, cells were treated with phospholipase C (PLC), an enzyme that cleaves GPI-anchored proteins. PLC (100 and 200 U/μL) significantly reduced ACE2-GPI surface staining and had no impact on ACE2 (Fig. 2D).

**Figure 2.**
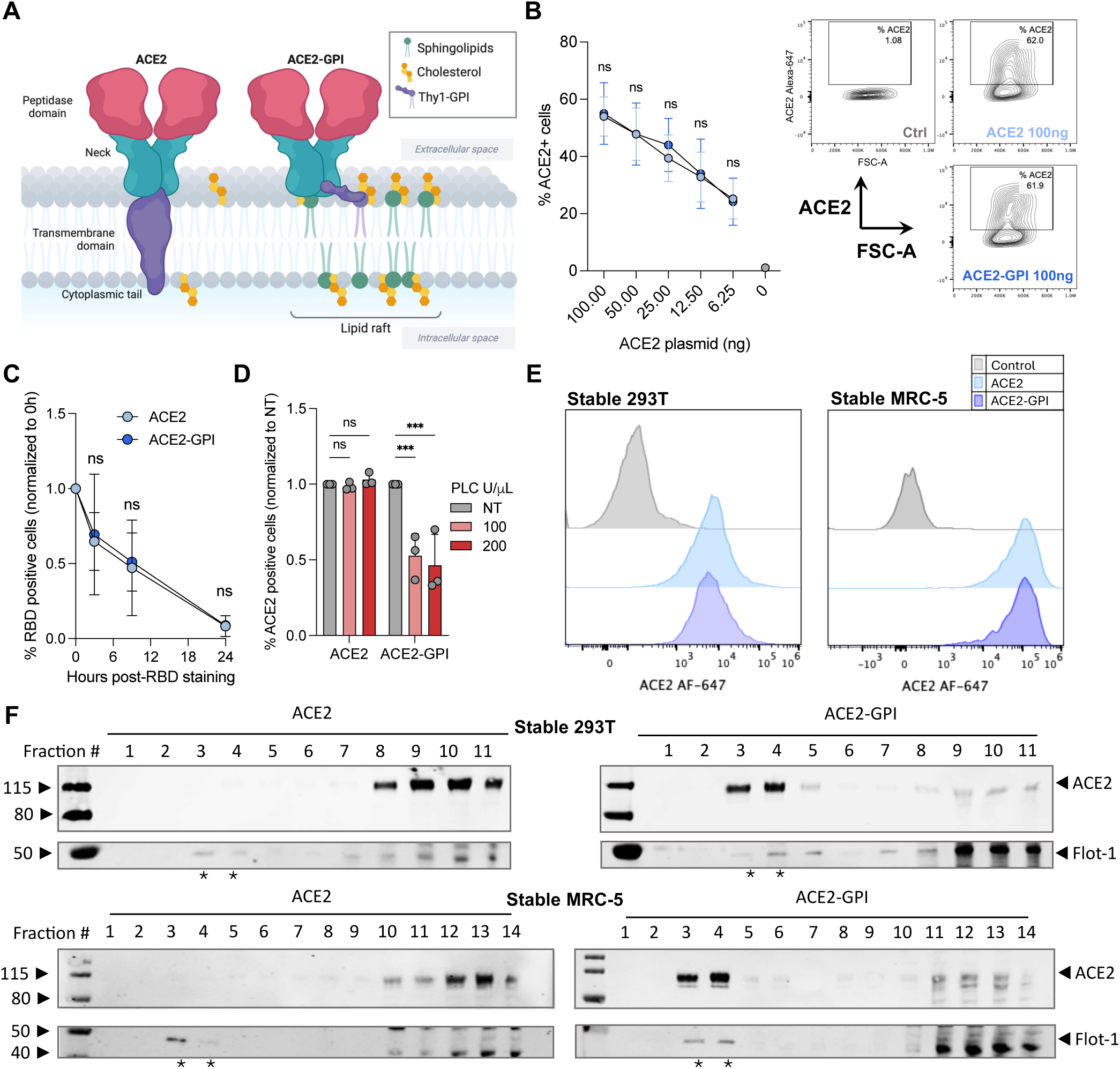
ACE2-GPI construct cell surface expression and lipid raft localization. **(A)** Schematic of the ACE2 and ACE2-GPI constructs used in this study. For ACE2-GPI, the transmembrane domain and cytoplasmic tail of ACE2 were replaced with Thy1, a GPI-anchored protein, to localize ACE2 to lipid rafts. **(B)** Surface expression of ACE2 and ACE2-GPI by flow cytometry analysis by ACE2 surface staining with anti-ACE2 VHH-B07-Fc. A serial dilution of ACE2-expression vector was used to transfect 293T cells before surface staining. Shown is the total percentage of ACE2-positive cells (left) and representative contour plots taken from one independent repeat (right). Data are means ±s.d. of three independent experiments. **(C)** 293T cells expressing ACE2 or ACE2-GPI were incubated with recombinant biotinylayed-SARS-CoV-2 RBD (Wuhan variant). Cell surface staining with streptavidin AF-488 was carried out at indicated timepoints. Data were normalized to 0h post-RBD staining. Data are means ±s.d. of three independent experiments. **(D)** Treatment of 293T cells with phospholipase C at indicated concentrations for 2 hours at 37°C. ACE2 surface levels were assessed by flow cytometry. Data were normalized to NT conditions. Data are means ±s.d. of three independent experiments. **(E)** Histogram plots from flow cytometry analysis of ACE2 cell surface staining of 293T cells (left) and MRC-5 cells (right) stably expressing ACE2 or ACE2-GPI. Results are representative of at least three independent experiments. **(F)** Cell lysates of 293T cells (above) and MRC-5 cells (below) stably expressing ACE2 or ACE2-GPI were ultracentrifuged in an iodixanol gradient to isolate membranes on their lipid content. * = Lipid raft associated protein, flotillin-1, was used as a control for lipid raft localization. For B, C, and D, Mann-Whitney tests were performed to compare ACE2 and ACE2-GPI conditions, \**P* < 0.05, \*\**P* < 0.001, \*\*\**P* < 0.0001, ns = not significant.

To investigate ACE2 membrane domain localization, we generated HEK 293T and MRC5, stably expressing ACE2 or ACE2-GPI cell lines. Both cell lines showed comparable surface levels of ACE2 and ACE2-GPI by flow cytometry (Fig. 2E). To assess localization, we used a membrane flotation assay. Flotation assays use an iodixanol gradient to separate membranes based on their lipid content (Vogt & Ott, 2015). Cells are lysed mechanically to preserve the membranes and lysate is loaded at the bottom of the gradient. During ultracentrifugation, membranes with higher lipid content, such as lipid rafts, will move up the gradient. Separation of the cell membranes revealed in both the HEK 293T and MRC5 stable cell systems, ACE2 localized to non-raft fractions whereas ACE2-GPI primarily localized to raft fractions as confirmed by flotillin-1 staining (Fig. 2F). A similar raft profile was observed in HEK 293T cells transiently transfected with ACE2 or ACE2-GPI (Fig. S1). Thus, ACE2 was non-raft-associated in the cell lines tested while ACE2-GPI localized to the raft compartment. Both proteins displayed similar surface expression and interaction with SARS-CoV-2 RBD.

### ACE2 and ACE2-GPI expressing cells display similar levels of SARS-CoV-2 pseudovirus entry and SARS-CoV-2 virus-induced fusion and replication

Localization of ACE2 to lipid rafts is suggested to enhance SARS-CoV-2 entry. To investigate this, we infected transfected HEK 293T and stable MRC5 cells with SARS-CoV-2 pseudovirus (Fig 3A). HEK 293T cells were co-transfected with TMPRSS2 and ACE2 or ACE2-GPI and infected with two doses of SARS-CoV-2 D614G pseudovirus expressing luciferase. The luciferase signal (RLU) correlated with the viral inoculum. Both ACE2 and ACE2-GPI HEK 293T cells showed no significant dìerences in the mean RLU in the absence of TMPRSS2 (Fig. 3B). However, in the presence of TMPRSS2, viral entry was increased in the ACE2-GPI condition by 2.5-fold compared to ACE2, regardless of the viral inoculum (Fig. 3B). Conversely, Spike mediated pseudovirus entry in ACE2 or ACE2-GPI stable MRC5 cells showed no significant dìerence when overexpressing ACE2 or ACE2-GPI (Fig. 3C).

**Figure 3.**
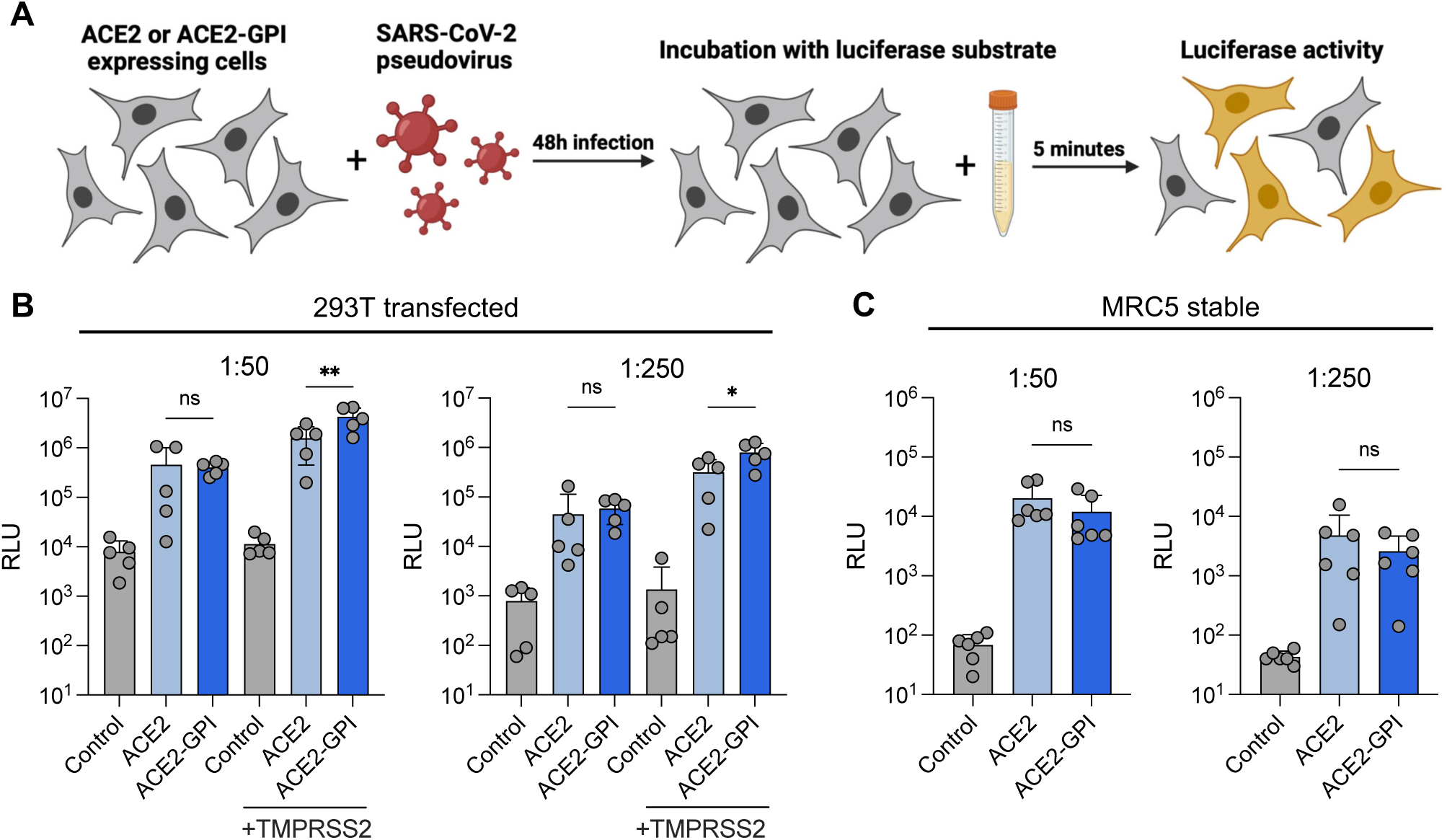
SARS-CoV-2 pseudovirus exhibits similar entry in ACE2 and ACE2-GPI expressing cells. **(A)** Schematic of experimental workflow. **(B)** 293T cells were transfected with ACE2 or ACE2-GPI 24 hours prior to pseudovirus infection. Infection was performed at a dilution of 1:50 (left) or 1:250 (right) of the virus for 48 hours before addition of luciferase substrate and measurement of luciferase activity. Data are means ±s.d. of five independent experiments. **(C)** As shown in B but using MRC5 stably expressing ACE2 or ACE2-GPI cells. Data are means ±s.d. of six independent experiments. For B and C, Mann-Whitney tests were performed to compare ACE2 and ACE2-GPI conditions, \**P* < 0.05, \*\**P* < 0.001, ns = not significant.

We next explored the impact of ACE2 localization on ancestral and post omicron SARS-CoV-2 variant entry and replication. Transfected HEK 293T GFP-Split and stable MRC5 GFP-split cells were infected with D614G and XBB.1.5 strains, 48h post infection syncytia formation was assessed by image quantification of GFP-Split complementation and replication was assessed by qPCR in the supernatant (Fig. 4A). As we have previously shown (Buchrieser et al., 2020; Planas et al., 2021), GFP positive syncytia were also positive for Spike and thus indicated productive SARS-CoV-2 infection (Fig. 4B). GFP area quantification showed no significant dìerences in D614G or XBB.1.5 variant induced syncytia formation by ACE2 or ACE2-GPI in HEK 293T and MRC5 cells (Fig. 4C & 4D). While the HEK 293T cells did not permit èicient replication, there was a non-significant increase in viral RNA copies/mL in ACE2-GPI expressing cells for both D614G and XBB.1.5 with a 2.9-fold and 1.5-fold increase respectively (Fig. 4E). In accordance with the prior pseudovirus entry results, the localization of ACE2 had no significant impact on viral replication in MRC5 cells for either D614G or XBB.1.5 (Fig. 4F). Overall, ACE2 localization did not greatly impact SARS-CoV-2 replication and syncytia formation.

**Figure 4.**
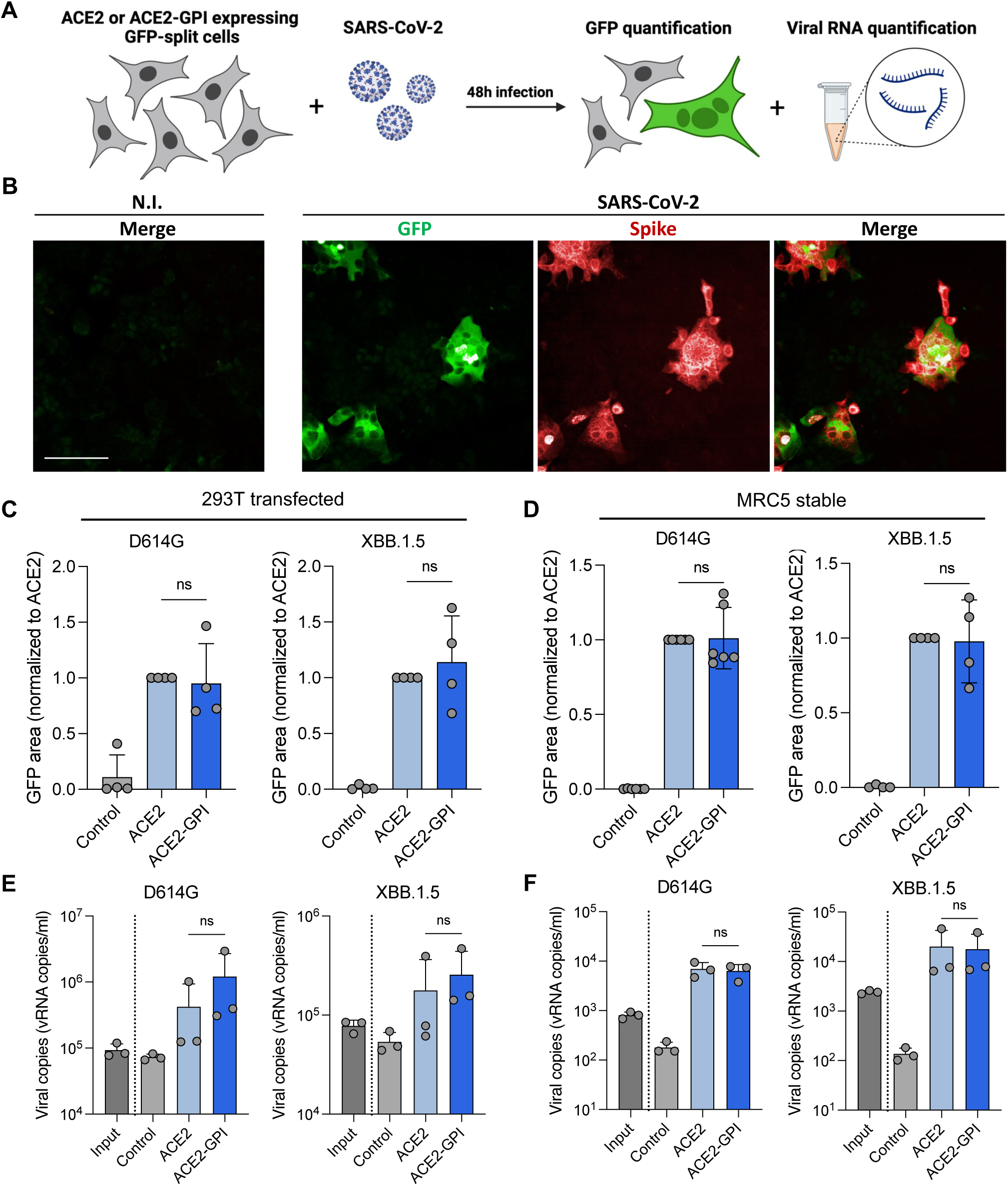
SARS-CoV-2 virus displays similar levels of fusion and replication in ACE2 and ACE2-GPI expressing cells. **(A)** Schematic of experimental workflow. **(B)** Immunofluorescence images of MRC5 GFP-split cells expressing ACE2 infected with SARS-CoV-2 for 48 hours and stained for Spike. Scale bar: 100 μm. **(C)** 293T GFP1-10 and GFP11 cells were co-transfected for 24 hours with ACE2 or ACE2-GPI before infection with SARS-CoV-2 D614G or XBB.1.5 variants at MOI 0.1 and 0.01. 48 hours post infection (hpi) cells were fixed and GFP area was quantified as a readout of fusion. Data were normalized to D614G fusion with ACE2. Data are means ±s.d. of four independent experiments. **(D)** As shown in C but with MRC5 cells stably expressing ACE2 or ACE2-GPI. Data were normalized to D614G fusion with ACE2. Data are means ±s.d. of four or five independent experiments. **(E)** Viral RNA quantification by qRT-PCR from the cell supernatant of infected 293T cells taken 48 hpi. Primers targeted the SARS-CoV-2 nucleoprotein. Data are means ±s.d. of three independent experiments. **(F)** Viral RNA quantification by qRT-PCR from the cell supernatant of infected MRC5 cells taken 48 hpi. Primers targeted the SARS-CoV-2 nucleoprotein. Data are means ±s.d. of three independent experiments. For C, D, E, and F, Mann-Whitney tests were performed to compare ACE2 and ACE2-GPI conditions, \**P* < 0.05, \*\**P* < 0.001, ns = not significant.

### ACE2 raft localization does not impact Spike-mediated syncytia formation

To further verify the impact of ACE2 localization on SARS-CoV-2 fusion, we examined syncytia formation through expression of Spike alone in HEK 293T and MRC5 cells. HEK 293T GFP-Split were transfected with Spike (D614G or XBB.1.5), ACE2 or ACE2-GPI and TMPRSS2 and fusion was assessed by GFP area quantification (Fig. 5A). In HEK 293T cells, ACE2-GPI significantly increased fusion compared to ACE2 by 1.5-fold and 3.0-fold with D614G and XBB.1.5 Spikes, respectively (Fig. 5B). In the presence of TMPRSS2, this increase with ACE2-GPI was less marked however the trend remained. Surface expression of ACE2 and ACE2-GPI was similar, and the levels of fusion were non-saturating (Fig. S2A). MRC5 cells are dìicult to transfect, we thus transfected HEK 293T GFP11 cells with Spike and co-cultured them stable ACE2 or ACE2-GPI MRC5 GFP1-10. Here, we found no significant dìerence in the level of Spike induced fusion with ACE2 or ACE2-GPI expressing cells for either D614G or XBB.1.5 Spike, in accordance with previous results (Fig. 5C & S2B).

**Figure 5.**
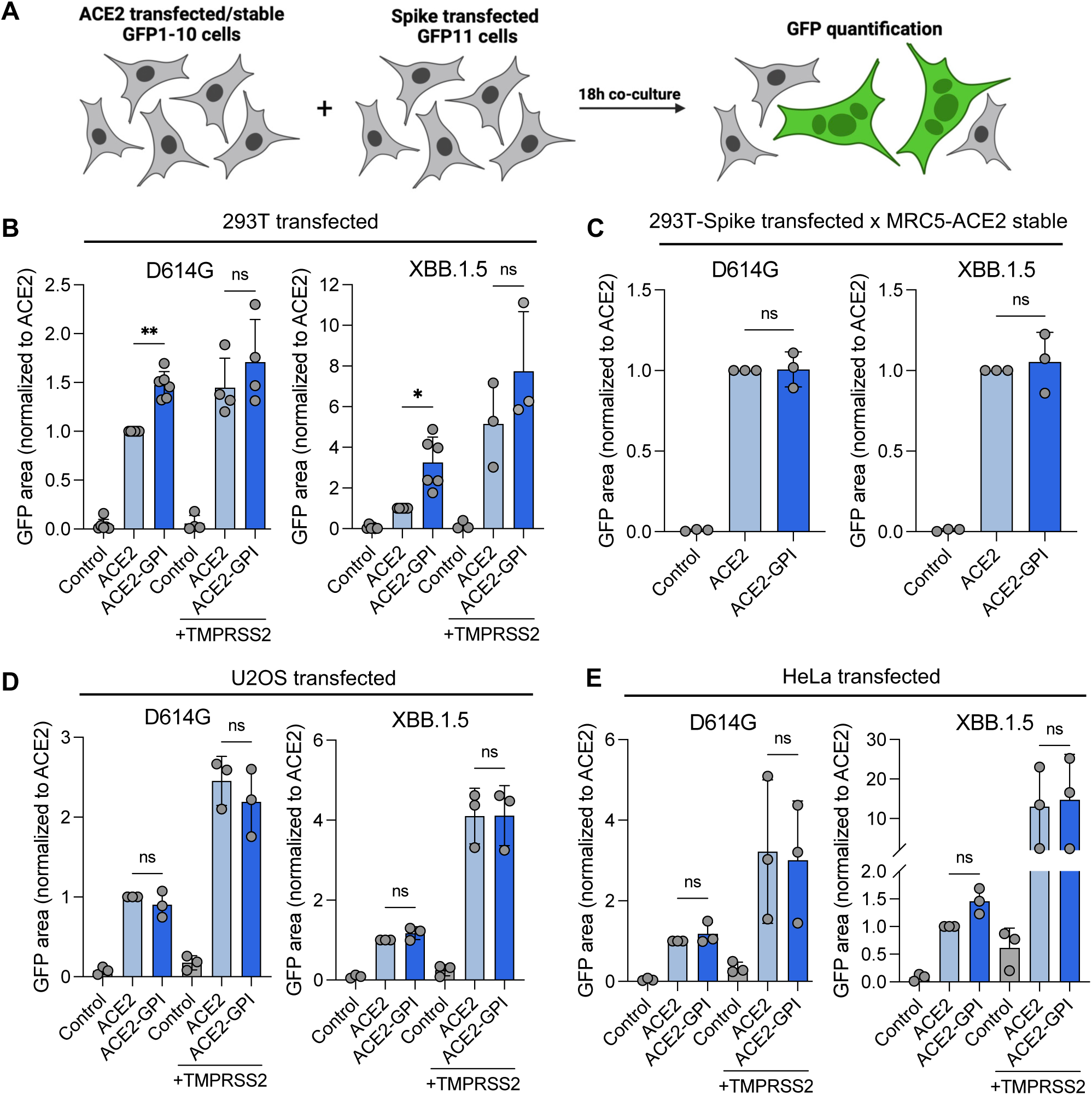
SARS-CoV-2 Spike exhibits similar levels of fusion with ACE2-GPI compared to ACE2. **(A)** Schematic of experimental workflow. **(B)** 293T GFP-split cells were transfected with SARS-CoV-2 D614G or XBB.1.5 Spikes and ACE2 or ACE2-GPI in the presence or absence of TMPRSS2 for 18 hours. Fusion was quantified by GFP area and data were normalized to D614G fusion with ACE2. Data are means ±s.d. of three, four, or five independent experiments. **(C)** 293T GFP 1-10 cells were transfected with Spike and co-cultured with MRC5 cells stably expressing ACE2 or ACE2-GPI for 18 hours. Fusion was quantified by area of GFP, and data were normalized to D614G fusion with ACE2. Data are means ±s.d. of three independent experiments. **(D)** As shown in B but using U2OS GFP-split expressing cells. Data were normalized to D614G fusion with ACE2. Data are means ±s.d. of three independent experiments. **(E)** As shown in B but using HeLa GFP-split expressing cells. Data were normalized to D614G fusion with ACE2. Data are means ±s.d. of three independent experiments. For B, C, D, and E, Mann-Whitney tests were performed to compare ACE2 and ACE2-GPI conditions, \**P* < 0.05, \*\**P* < 0.001, ns = not significant.

To further clarify the impact of ACE2 localization on Spike-induced syncytia formation, we carried out the same fusion assay as with HEK 293T cells in two additional cell lines, U2OS and HeLa cells. Here, for both cell lines we saw no dìerence in the levels of fusion between ACE2 and ACE2-GPI with both D614G and XBB.1.5 Spikes (Fig. 5D & 5E). Levels of fusion were non-saturating and ACE2 and ACE2-GPI surface expression were not significantly dìerent (Fig. S2C & S2D). While TMPRSS2 increased the overall levels of fusion there were no significant dìerences between ACE2 and ACE2-GPI fusion. Altogether, the localization of ACE2 into lipid rafts had little impact on Spike-induced fusion in dìerent cell lines, including U2OS, HeLa and MRC5 lung fibroblast cells, except for HEK 293T cells where we saw a trend to increase fusion with raft localization, a finding that requires further exploration.

## DISCUSSION

We describe the role of cholesterol and ACE2 lipid raft localization during SARS-CoV-2 fusion. Cholesterol is required for membrane fusion of various enveloped viruses including lentiviruses, flaviviruses, and coronaviruses, consistent with our results (Osuna-Ramos et al., 2018; Roncato et al., 2022; Salimi et al., 2020). Increasing membrane cholesterol by media supplementation increases SARS-CoV-2 pseudovirus entry while cholesterol depletion using agents such as MβCD, statins, or cholesterol 25-hydroxylase has the reverse èect (El Khoury & Naim, 2023; X. Li et al., 2021; Wang et al., 2020; Zang et al., 2020). This is in line with clinical reports describing the reduction of COVID-19 disease severity by statins (Kouhpeikar et al., 2022; REMAP-CAP Investigators, 2023). Cholesterol is required in membrane fusion due to its intrinsic ability to stabilize curved fusion intermediates and facilitate the penetration of fusion peptides into the target membrane (Yang et al., 2016). While cholesterol is a key component of lipid rafts, its depletion is not specific to rafts despite MβCD often being referenced as a lipid raft inhibitor (Edidin, 2003; Mahammad & Parmryd, 2008). Thus, cholesterol depletion may disturb various cellular functions and membrane properties, notably those involved in viral fusion.

To assess the role of ACE2 localization without cholesterol depletion and lipid raft disruption we design an ACE2-GPI construct. This led to re-localization of ACE2 from non-lipid raft regions to lipid raft regions. The GPI anchor is a post-translational modification comprised of a phosphoethanolamine linker attached to a glycan core and phospholipid tail that preferentially insert into the outer leaflet of lipid rafts (Paulick & Bertozzi, 2008). Attaching a GPI anchor to angiotensin-converting enzyme (ACE) sequesters the protein to lipid rafts while wild-type ACE does not and treatment of the cells with PLC, releases the protein from the surface (Parkin et al., 2003). Furthermore, the cytoplasmic tail of ACE2 is not required for either SARS-CoV or SARS-CoV-2 entry and should thus not impact the function of ACE2-GPI as a receptor (Karthika et al., 2021).

Through membrane flotation assays we show that ACE2 localized to non-raft membrane regions while ACE2-GPI localized preferentially within lipid rafts. ACE2 has been previously described as a lipid-raft associated protein in studies exploring SARS-CoV and SARS-CoV-2 entry in VeroE6 and Caco2 cell lines (El Khoury & Naim, 2023; X. Li et al., 2021). Conversely, other studies in VeroE6 and CHO cells conclude that ACE and ACE2 are found predominantly in the detergent-soluble membrane domains, separate to the lipid rafts (G. M. Li et al., 2007; Warner et al., 2005). From our results, we postulate that ACE2 is a non-raft associated protein in the human cell lines that we tested. This warrants further investigation including the localization of ACE2 in primary human cells and tissue which remains unexplored.

We describe little èect of ACE2 raft localization on SARS-CoV-2 entry, or virus and Spike induced-syncytia formation in several cell lines. HEK 293T cells expressing ACE2-GPI exhibited a slight but general increase in these processes compared to ACE2 expressing cells, an observation that requires further research. Regarding viral entry, internalization of receptors in rafts occurs via caveolae-mediated endocytosis as opposed to the classical clathrin-dependent pathway (Roncato et al., 2022). While several studies suggest ACE2-raft localization is important for SARS-CoV-2 entry and fusion, SARS-CoV-2 utilises both endosomal pathways to mediate entry into the host cell, together with cell surface fusion, suggesting ACE2 raft localization is not essential for this process (Bayati et al., 2021; Jackson et al., 2022).

Palmitoylation is a post-translational modification of cysteine residues within the CT of envelope proteins including SARS-CoV-2 Spike (Wu et al., 2021), influneza haemagglutinin (Chlanda et al., 2016), MLV env (M. Li et al., 2002), and HIV-1 gp160 (Rousso et al., 2000). The addition of fatty acids allows anchoring of the envelope proteins to the plasma membrane and mutation of these residues strongly deceases viral infectivity. Additionally, aromatic SARS-CoV Spike residues at the membrane interface facilitate interaction with membrane cholesterol (Corver et al., 2009; Meher et al., 2019). As with ACE2, Spike requires cholesterol for fusion and this mechanism is independent of lipid rafts (Sanders et al., 2021). Thus, we propose a model by that, while cholesterol in the plasma membrane is required for SARS-CoV-2 fusion, the localization of ACE2 into cholesterol-rich lipid rafts does not enhance this process.

<u>T</u>his study contains limitations. First, due to the artificial nature of the ACE2-GPI construct, we rely on overexpression systems and cell-line based models to assess viral entry and syncytia formation. Further research may look to more physiologically relevant models, for example, expressing ACE2-GPI in primary epithelia air-liquid interface models. Additionally, we do not factor the localization of other proteins required for SARS-CoV-2 fusion such as proteases including TMPRSS2 and cathepsins. Finally, we demonstrate ACE2 localization through membrane flotation assays however this would be supported by techniques such as fluorescence microscopy through co-localization analysis.

Overall, the data presented here show the requirement of cholesterol to facilitate entry of SARS-CoV-2 as well as virus and Spike-mediated fusion. ACE2 localizes outside of lipid rafts in the cell line models we tested. Re-localization of ACE2 to lipid rafts had little or no impact on viral entry and fusion. The independence of ACE2 lipid raft localization provides insight into the mechanisms of SARS-CoV-2 entry and fusion.

## ACKNOWLEDGEMENTS

The authors thank the members of the Virus and Immunity Unit (Institut Pasteur) for their discussion and help and the stà at the UtechS Photonic BioImaging core facility (Institut Pasteur). W.B. was supported by a stipend from the Pasteur-Paris University (PPU) International PhD program.

## FUNDING

Work in the UVI unit is funded by Institut Pasteur, Urgence COVID-19 Fundraising Campaign of Institut Pasteur, Fondation pour la Recherche Médicale, ANRS-MIE, the Vaccine Research Institute (ANR-10-LABX-77), Labex IBEID (ANR-10-LABX-62-IBEID), ANE/FRM Flash COVID PROTEO-SARS-CoV-2, ANR Coronamito, the HERA Project DURABLE (grant 101102733) and the HERA project LEAPS. The Opera Phenix system was co-funded by Institut Pasteur and the Région Ile de France (DIM1Health). The funders of this study had no role in study design, data collection, analysis, interpretation, or writing of the article.

## AUTHOR CONTRIBUTIONS

Conceptualization: O.S., J.B., W.B., N.C.

Methodology: W.B., J.B., O.S., I.M., N.C., F.G-B., F.P.

Investigation: W.B., J.B., O.S., I.M., N.C.

Data collection and analysis: W.B., I.M., C.P., J.B.

Manuscript writing and editing: W.B., J.B., O.S.

## DELCLARATION OF INTEREST

The authors declare no competing interests

**Supplementary figure 1.** Cell lysates of transfected 293T cells expressing ACE2 or ACE2-GPI were ultracentrifuged in an iodixanol gradient to isolate lipid raft and non-lipid raft membrane fractions. * = Lipid raft associated protein, flotillin-1, was used as a control for lipid raft localization.

**Supplementary figure 2.** Cell surface expression controls measured by flow cytometry for ACE2, Spike and TMPRSS2 together with raw image data showing GFP-positive syncytia in 293T cells **(A)**, U2OS cells **(B)**, HeLa cells **(C)**, and MRC5 cells **(D)** taken from one independent replicate of each. Scale bars for A and C = 400 μm. Scale bars for B and D = 200 μm. For all panels Mann-Whitney tests were performed to compare ACE2 and ACE2-GPI conditions, ns = not significant.

